# Selective degradation of platelet BTK by PROTAC NX-5948 provides antithrombotic benefits without affecting haemostasis

**DOI:** 10.64898/2025.12.02.691893

**Authors:** Carl J. May, Justin S. Trory, Connor E. Webb, Hetty S. Walker, Yong Li, Alastair W. Poole, Jordan Vautrinot, Jay Tromans, Ingeborg Hers

## Abstract

Current antithrombotic therapies are effective in reducing thrombotic events but are limited by their associated risk of bleeding. BTK acts as a key signalling switch that drives platelet activation during thrombosis but is largely dispensable for routine haemostasis. It is an important non-redundant signalling mediator downstream of the GPVI and CLEC-2 receptors, plays a key role in thrombosis with minimal involvement in haemostasis, making it an attractive antithrombotic target. While BTK inhibitors effectively reduce thrombosis, their clinical use has been limited due to off-target effects. Protein degraders may overcome this limitation by enabling the ubiquitin proteasomal system to selectively target and degrade BTK. We here assessed the ability of the BTK degraders NX-2127 and NX-5948, currently in clinical trials for B cell pathologies, to target platelet BTK for degradation. NX-2127 and NX-5948 induced concentration-dependent degradation of BTK in washed platelets, platelet-rich plasma and whole blood. NX-5948 showed no hook effect and outperformed NX-2127 in potency, efficacy, and degradation kinetics. TMT proteomic analysis confirmed selective BTK degradation by NX-5948 with no evidence of major off-target effects. BTK degradation impaired CRP-mediated integrin αIIbβ_3_ activation, P-selectin expression, platelet aggregation and *in vitro* thrombosis, with PAR-1 mediated platelet function being left intact. Dosing mice with NX-5948 led to efficacious degradation of platelet BTK and impaired CRP-, but not thrombin-, mediated *ex vivo* platelet function. *In vivo*, arterial thrombosis was markedly reduced, without an increase in bleeding time. Together, these results highlight NX-5948 as a potent, selective BTK degrader with antithrombotic potential and minimal haemostatic impact.

**Key Points:**

1. The BTK degrader NX-5948 potently and selectively degrades platelet BTK and suppresses thrombus formation without affecting bleeding.
2. Targeting BTK degradation offers a new antithrombotic strategy that spares haemostasis and may aid patients intolerant to DAPT.

## Introduction

The abnormal formation of thrombi can lead to severe and potentially life-threatening consequences. By obstructing blood vessels and impairing circulation, thrombi can precipitate conditions such as myocardial infarction, pulmonary embolism, ischaemic stroke, and deep vein thrombosis^1^. Dual antiplatelet therapy (DAPT) with aspirin and a P2Y_12_ antagonist effectively reduces recurrent thrombotic events but also increases bleeding risk, limiting clinical use and highlighting the need for antithrombotic therapies that prevent pathological thrombosis without impairing haemostasis.

The collagen receptor glycoprotein VI (GPVI) and its downstream effector Bruton’s Tyrosine Kinase (BTK) have been proposed as promising antithrombotic targets, as they are critical contributors to thrombosis but are largely redundant for haemostasis. Their role is supported by impaired thrombus formation on collagen and protection from arterial thrombosis in GPVI-deficient models and humans with minimal bleeding^2–8^. BTK inhibitors impair atherosclerotic plaque induced thrombus formation in humans^9–11^ and reduce venous thrombosis in mice^12^. However, their therapeutic window remains narrow due to off-target bleeding^13,14^. Interestingly, patients with X-linked agammaglobulinemia (XLA), who lack functional BTK, do not exhibit an increased risk of bleeding, suggesting that selective inhibition of BTK is unlikely to affect haemostasis^15,16^.

To address the limitations of BTK inhibitors, such as their narrow therapeutic windows and off-target effects, BTK degraders present a promising new avenue for anti-thrombotic treatments, with the added benefit of eliminating both the kinase and scaffolding functions of BTK^17^. Over the last few years Proteolysis Targeting Chimeras (PROTACs) have shown great promise as targeted protein degraders. These bifunctional molecules recruit an E3 ligase to ubiquitinate and degrade the target via the Ubiquitin-Proteasomal System^18^. We previously demonstrated that human platelets are highly susceptible to PROTAC-mediated BTK degradation using the BTK degraders DD-03-171 and DD-04-15, with TMT-based proteomic analysis confirming superb specificity^19^.

As BTK also plays a central role in B-cell receptor signalling and sustained activation of BTK is a defining feature of many B-cell malignancies, significant industrial effort has been directed towards developing BTK PROTACs for the treatment of B-cell lymphomas. Several of these, including NX-2127 and NX-5948 are currently undergoing early-phase clinical trials. In addition to degrading BTK, NX-2127 also acts as a molecular glue resulting in degradation of the transcription factors, Ikaros (IKZF1) and Aiolos (IKZF3)^20^, which enhances the anti-tumour properties of T-cells^21^. NX-5948 is a related compound developed for similar indications but selectively targets BTK without affecting transcription factors^22^.

At present, the prospective application of these PROTACs as anti-thrombotic agents by targeting platelets remains to be explored. Given their potential to preserve haemostasis, we investigated the effects of NX-2127 and NX-5948 on platelets.

Specifically, we sought to characterise the ability of these clinical-stage BTK degraders to eliminate BTK from human and mouse platelets, to demonstrate their mechanistic basis and to assess the functional consequences on platelet activation, aggregation, thrombus formation and haemostasis.

## Methods

### Human Platelet Isolation

Blood from healthy, drug-free volunteers was collected into 4% (w/v) trisodium citrate following NHS Research Ethics Committee approval (20/SC/0222) and in accordance with the Declaration of Helsinki. Platelets were isolated as described previously. Briefly, blood was acidified with Acid Citrate Dextrose (ACD; 1:7, v/v), centrifuged at 150 × g for 17 min at room temperature (RT), and platelet-rich plasma (PRP) collected. Platelets were pelleted at 800 × g for 10 min in the presence of 2 µM prostaglandin E₁ (PGE₁) and 0.02 U/ml apyrase, washed in CGS buffer (13 mM trisodium citrate, 22 mM glucose, 120 mM NaCl) containing apyrase, and resuspended in HEPES-Tyrode’s buffer (145 mM NaCl, 3 mM KCl, 0.5 mM Na₂HPO₄, 1 mM MgSO₄, 10 mM HEPES, pH 7.4) supplemented with 0.1% (∼5.5 mM) D-glucose and apyrase to yield 4 × 10⁸ platelets/ml.

### Incubation with PROTACs

Platelets in whole blood, PRP, or washed suspensions were incubated at 30 °C with NX-2127 (HY-153220, MedChemExpress) or NX-5948 (8856, Tocris Bioscience) at the indicated concentrations and durations, then isolated as described above for human or mouse platelets.

### Western Blotting

Washed platelets were lysed in 4× NuPAGE sample buffer containing 0.1% (w/v) SDS. Lysates were analysed by SDS-PAGE and immunoblotting using primary antibodies against Talin (TA205, MA528133, Thermo Fisher/Invitrogen), GAPDH (6C5, SC-32233 Santa Cruz), BTK (D3H5, 8547, Cell Signaling Technology), and TEC (A12533, antibodies.com), as described previously^19^.

### In Vitro Thrombus Formation Assay

PRP was incubated for 4 hours with vehicle (DMSO), 3 µM NX-2127, or 100 nM NX-5948, then labelled with 2 µM DiOC₆ (3,3ʹ-dihexyloxacarbocyanine iodide) for 30 min. PRP was recombined with erythrocytes and perfused through Vena8™ GCS biochips coated with 50 µg/ml fibrillar collagen at 3.07 ml/h (shear rate = 1000 s⁻¹) for 5 min. Thrombus formation was imaged by confocal microscopy (two fields per channel). Images were analysed using a custom ImageJ plugin; data were processed in Microsoft Excel and statistics performed in GraphPad Prism.

### Platelet Spreading Assay

Washed platelets (5 × 10⁶/ml) were incubated for 75 min at 30 °C on coverslips coated with fibrinogen (20 µg/ml) and collagen (10 µg/ml). Platelets were fixed with 8% paraformaldehyde, permeabilised with 0.1% Triton X-100, and stained with ActinGreen™ 488 ReadyProbes™ (Invitrogen). Samples were mounted in ProLong™ Diamond Antifade Mountant and imaged using an EVOS-FL microscope (40× objective). Five fields per coverslip were captured per condition. Platelets were categorised into three spreading stages (Stage 1, resting; Stage 2, filopodia; Stage 3, lamellipodia/actin membrane) using a macro developed by Stephen Cross (University of Bristol). Data were analysed in Excel and plotted in GraphPad Prism.

### Mice

Male and female C57BL/6 mice (9-10 weeks; Charles River) were used under UK Home Office Project Licence PP5643338. All procedures were approved by the University of Bristol Animal Welfare and Ethical Review Body and conducted in accordance with the Animals (Scientific Procedures) Act 1986. Mice were randomised to receive NX-5948 (30 mg/kg, oral gavage) or vehicle control (n = 3-5 per group). Each animal constituted an independent experimental unit. Sample size was informed by prior thrombosis studies providing 80% power to detect ≥ 30% differences. Imaging and analysis were performed blinded to treatment. Mice were euthanised by rising CO₂ inhalation 24 hours post-treatment, and blood collected by cardiac puncture into 4% trisodium citrate (1:9, v/v).

### Mouse Platelet Isolation

Blood was collected by cardiac puncture using a 23 G needle pre-loaded with ACD. Samples were diluted with HEPES-Tyrode’s buffer containing glucose (HT+G) and 10 mM EGTA to 3 ml. PRP was obtained by centrifugation at 200 × g for 5 min (RT). The red cell layer was resuspended in HT+G + EGTA, centrifuged again, and the pooled PRP collected. Combined PRP was centrifuged with 0.5 µM PGE₁ (and 0.2 U/ml apyrase if required) at 800 × g for 10 min. The resulting pellet was washed once in HT+G with PGE₁, recentrifuged, and resuspended in a small volume of HT+G. Platelet concentration was measured and adjusted prior to use.

### In Vivo Arterial Thrombosis and Tail-Bleeding Assays

Mice were anaesthetised with ketamine (100 mg/kg; Vetalar V, Pfizer) and xylazine (10 mg/kg; Rompun, Bayer). Platelets were labelled intravenously with DyLight-488-anti-GPIbβ (100 µg/kg; X488, Emfret) 10 min before application of 12% ferric chloride to the carotid adventitia for 3 min. Fluorescence images were acquired for 20 min and analysed in ImageJ (NIH) as integrated thrombus fluorescence after background subtraction. For tail bleeding assays bleeding time was assessed using a warm-saline immersion method to minimise vasoconstriction and provide standardised physiological conditions. Mice (C57BL/6) were anaesthetised with ketamine (100 mg/kg; Vetalar V, Pfizer) and xylazine (10 mg/kg; Rompun, Bayer) and maintained on a heating pad to prevent hypothermia. A full-thickness transverse incision was made 5 mm from the tip using a fresh scalpel blade. Immediately after incision, the tail was immersed 1-2 cm into sterile saline pre-warmed to 37 °C in a 50 mL tube. Bleeding was monitored continuously, and the time to first cessation of bleeding (defined as no visible blood for 30 s) was recorded. Any subsequent rebleeding events were documented, and total bleeding time was measured up to a 10-minute maximum.

### Flow Cytometry

Integrin α_IIb_β_3_ activation and α-granule secretion were assessed in washed human platelets (2 × 10⁷/ml) using PAC1-FITC and anti-CD62P-PE antibodies, and in washed mouse platelets using Jon/A-PE and WugE9-FITC. Platelets were stimulated with CRP-XL (5 µg/ml, Triple Helical Peptides), TRAP-6 (10 µM, Bachem), or thrombin (1 U/ml) for 10 min, fixed with 2% paraformaldehyde, and analysed on a BD Accuri™ C6 Plus flow cytometer (FL1/FL2 channels), capturing 10,000 events per sample. Results are presented as median fluorescence intensity (MFI).

### Calcium Mobilisation Assay

Calcium mobilisation was measured as described previously ^24^. Briefly, platelet-rich plasma (PRP) was prepared as outlined above and adjusted to 4 × 10⁸ platelets/ml in CGS buffer (13 mM trisodium citrate, 120 mM NaCl, 22 mM D-glucose, pH 6.5) containing 0.02 U/ml apyrase. Platelets were loaded with 4 µM Fura-2 AM in the presence of 0.35% fatty acid-free bovine serum albumin (BSA) for 30 min at 30 °C. After loading, samples were centrifuged at 520 × g for 10 min (RT) and resuspended in HEPES-Tyrode’s buffer (HT) supplemented with 0.02 U/ml apyrase and 5.5 mM D-glucose. Platelet counts were verified using a Coulter counter (Beckman Coulter Z1).

For agonist stimulation, partial half-log concentration-response curves were generated for CRP-XL and TRAP-6, with ionophore A23187 included as a positive control. Immediately before loading, washed platelets (4 × 10⁸/ml) were diluted 1:1 in HEPES-Tyrode’s buffer and re-calcified with 1 mM CaCl₂. Aliquots (95 µl; 2 × 10⁸/ml) were added to a 96-well clear-bottom black microplate (Corning, Fisher Scientific) and preincubated for 5 min at 37 °C. Intracellular Ca²⁺ mobilisation was recorded in real time using a Tecan Infinite 200 PRO multimode plate reader (Männedorf, Switzerland) at 340/380 nm excitation and 510 nm emission, with 1-s shaking intervals between reads. Data were expressed as the 340/380 nm fluorescence ratio.

### Proteomic Analysis of Platelet Samples

Washed platelets were lysed in 1× RIPA buffer (25 mM HEPES, 200 mM NaCl, 1 mM EDTA, 2% NP-40, 0.5% sodium deoxycholate, 0.1% SDS, pH 7.4) supplemented with PhosSTOP™ and cOmplete™ Mini EDTA-free inhibitors (Roche). Protein concentrations were determined by BCA assay (Pierce™, Thermo Fisher, Cat. 23225) and normalised. Proteins were digested with trypsin, TMTpro-labelled, pooled, and desalted using SepPak cartridges. Peptides were fractionated by high-pH reversed-phase chromatography into 20 fractions and analysed by nano-LC-MS/MS on an Orbitrap Fusion Lumos (SPS-MS³ mode). Data were searched against UniProt Human and contaminant databases (Sequest; 5% FDR). Log₂-transformed protein abundances were compared (PROTAC-DMSO) by paired t-test and visualised as volcano plots in R (ggplot2 v4.2.3). Single-peptide identifications were manually inspected to understand isoform assignments and investigated by western blotting. Apparent degradation of the adaptor-associated kinase 1 (AAK1) annotated protein group by NX-2127 was driven by a single unique peptide mapping to a poorly annotated non canonical AAK1-like isoform (A0A096LP25; 3% coverage) and no peptides uniquely identifying canonical AAK1 (Q9BUD9). Western blotting of NX-2127 treated platelets for AAK1 showed no evidence of AAK1 degradation (**Figure S2**).

### 3C-Well Plate Aggregation

Platelet aggregation was assessed in a 96-well format. PRP was prepared and incubated with PROTACs, then dispensed into half-area plates alongside platelet-poor plasma (PPP) controls representing 100% aggregation. Plates were warmed to 37 °C, stimulated with agonists, and shaken at 1200 rpm for 5 min. Absorbance (595 nm) was measured on a Labtech LT4500 reader. Percentage aggregation was calculated as the normalised absorbance of stimulated PRP minus that of unstimulated PRP, each relative to PPP (response-PPP).

### Statistical Analysis

Data were analysed in GraphPad Prism (v9). Welch’s t-test was used for two-group comparisons and one-way ANOVA for analyses involving three or more groups. Statistical significance was defined as p ≤ 0.05 (*), p ≤ 0.01 (**), p ≤ 0.001 (***), and p ≤ 0.0001 (****).

## Results

### NX-2127 and NX-5348 efficiently and selectively degrade BTK in human platelets

We examined whether the cereblon-recruiting BTK degraders NX-2127 and NX-5948 could eliminate BTK protein in human platelets, a cell type only recently shown to be susceptible to PROTAC-mediated targeted protein degradation ^19^. Concentration-response analyses in washed platelets and platelet-rich plasma (PRP) revealed that both compounds robustly degraded BTK, but with striking differences in potency and cooperativity.

In washed platelets, NX-2127 produced a concentration-dependent decline in BTK with a DC₅₀ of 12 nM and a classic “hook effect” at high concentrations, consistent with non-productive ternary complex formation when a PROTAC binds to either the E3 ligase or the target protein, but not both at the same time. (**Figure 1A**). In PRP, its potency was reduced roughly tenfold (DC₅₀ ≈ 98 nM; **Figure 1C**), suggesting partial sequestration by plasma proteins. NX-5948 achieved a DC₅₀ of 5.3 nM in washed platelets (**Figure 1B**) and maintained near-complete degradation without any hook effect, suggesting highly cooperative target engagement. The potency shift between washed platelets and PRP was modest (DC₅₀ = 12.8 nM; **Figure 1D**), implying that NX-5948 remains active even in plasma, possibly due to reduced susceptibility to protein binding than NX-2127. The relative resistance of NX-5948 to plasma sequestration represents a favourable pharmacological property for in vivo use. Comparable degradation in PRP and whole blood confirmed that NX-5948 was not sequestered within red cells (**Figure S1**).

**Figure 1:**
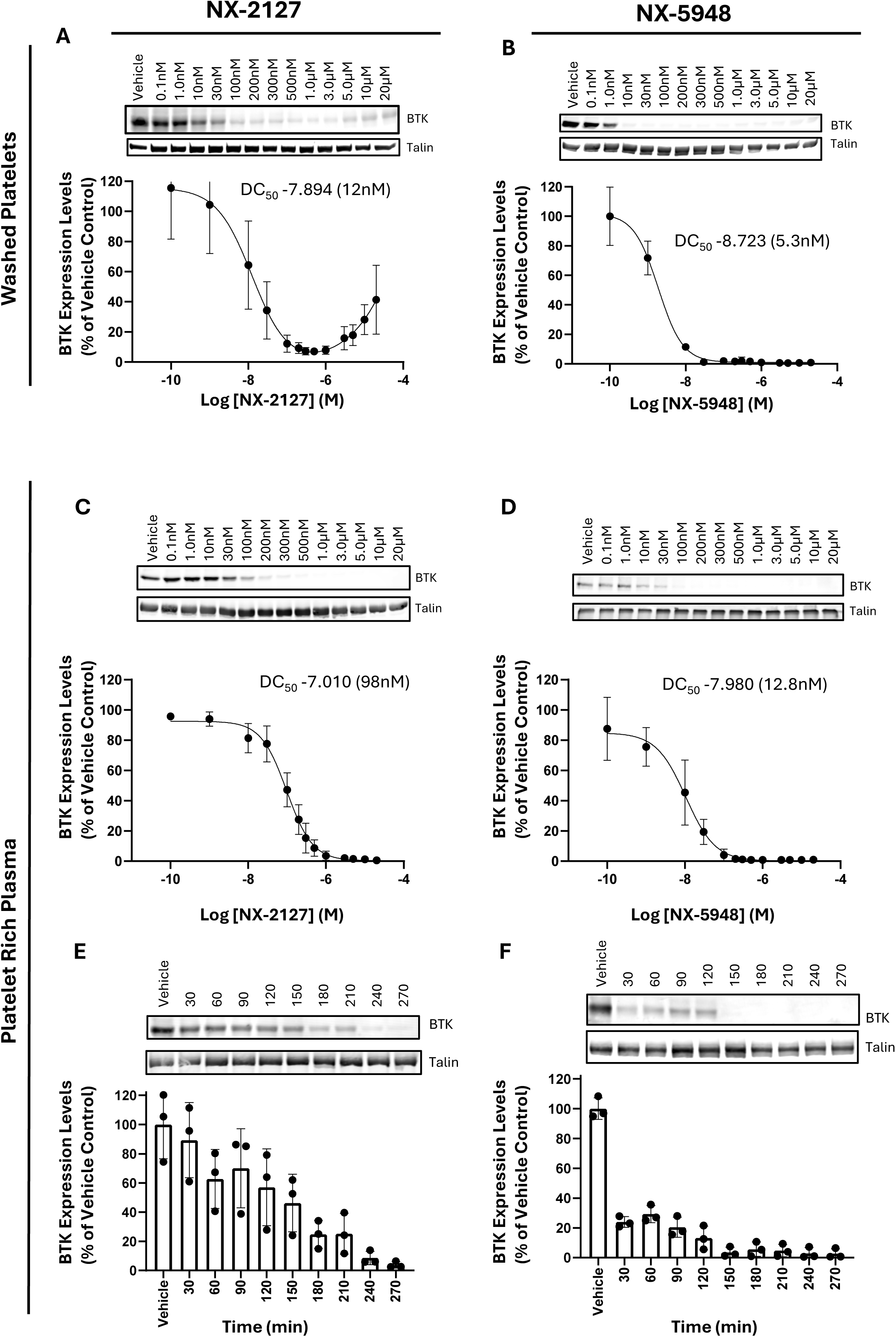
Concentration-dependent reduction of BTK expression by NX-2127 and NX-5948 in human platelets. (A-D) Concentration-response of NX-2127 and NX-5948 in washed human platelets (**A, B**) and platelet rich plasma (PRP, **C, D**). Platelets were treated with the indicated concentrations of NX-2127 (**A,C**) and NX-5948 (**B,D**) for 4 hours at 30 °C. BTK levels were quantified by densitometry of western blot and normalised to vehicle (DMSO) treated control. The DC_50_ in washed platelets for NX-2127 and NX-5948 was 12 nM and 5.3 nM, respectively (calculated using non-linear regression in GraphPad PRISM). The DC_50_ in PRP for NX-2127 and NX-5948 was 98 nM and 12.8 nM, respectively. **(E,F)** Time-dependent degradation of BTK by NX-2127 (**E**) and NX-5948 (**F**) in human PRP. Human platelets in PRP were treated with 3.0 μM of NX-2127 and 100 nM of NX-5948 for the indicated time points ranging from 30 min to 270 min. BTK levels were quantified by densitometry of western blot and normalised to vehicle (DMSO) treated control. The curves were generated from quantification of western blots (mean ± SEM, n=3 biological replicates) with representative blots shown above the respective curve, Talin used as a loading control for normalisation.

Kinetic degradation profiling reinforced these distinctions. NX-2127 required ∼150 min to reach 50% BTK loss and 250 min for maximal degradation, whereas NX-5948 achieved the same endpoints within 30 min and 150 min, respectively (**Figures 1E-F**). This rapid and near-complete loss of BTK establishes NX-5948 as a highly potent and kinetically efficient degrader.

### NX-2127 and NX-5348 have limited effect on TEC

Quantitative TMT-proteomics revealed high substrate specificity of the two degraders. Under matched conditions (3 µM NX-2127, 100 nM NX-5948), NX-2127 and NX-5948 induced degradation of BTK with high efficacy. (**Figure 2A-B**). Although a reduction in the level of non-canonical AAK1 (A0A096LP25) was detected following NX-2127 treatment, western blotting confirmed AAK1 expression levels were unaltered (Figure S2). Both compounds achieved near-total BTK loss, but NX-5948 did so at submicromolar concentrations (500 nM versus 20 µM for NX-2127), underscoring a > 40-fold pharmacological advantage. Neither degrader affected the related TEC kinase by proteomics or immunoblotting (**Figure 2C**). The narrow degradation profile of both NX-2127 and NX-5948 indicates that neither PROTAC induces the broader kinome destabilisation reported for some early BTK degraders ^25^. Mechanistic validation using pomalidomide or a NEDD-activating-enzyme inhibitor confirmed that BTK loss required CRBN-mediated ubiquitin-proteasomal activity (**Figure 3A-B**).

**Figure 2:**
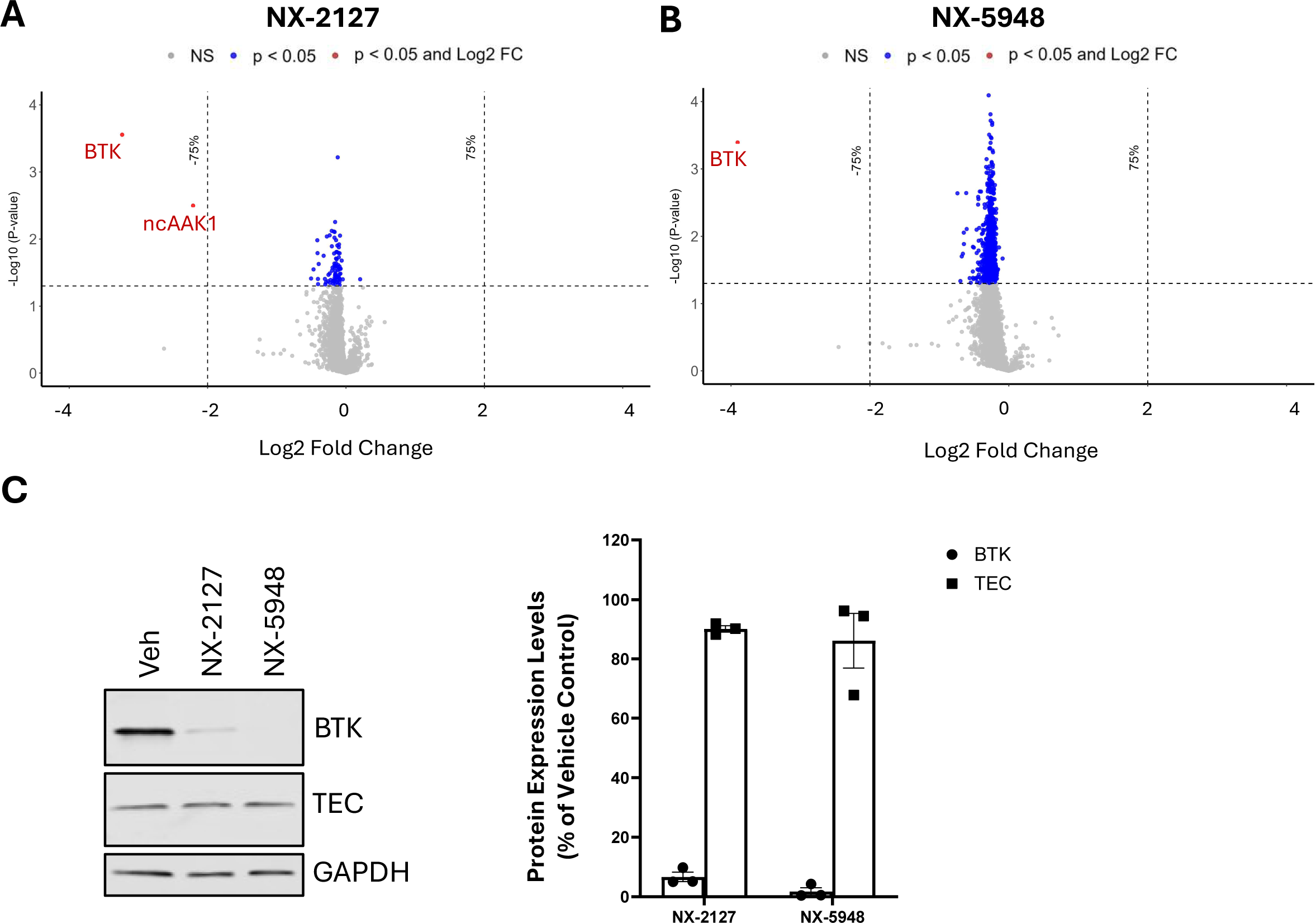
Specificity of degradation by NX-2127 and NX-5948 in human platelets (A-C) Human platelets in PRP were treated with 3.0 μM NX-2127 (A) or 100 nM NX-5948 (**B**) for 4 hours at 30°C before being washed, lysed and analysed by TMTpro-based quantitative proteomic analysis **(A**-**B)** or western blotting **(C)**. Proteins significantly downregulated (p < 0.05) are shown in red. Proteins exhibiting >75% (chosen a priori) reduction relative to vehicle-treated levels and meeting the same significance threshold are shown in blue (n=3 biological replicates) (**A,B**). NX-2127 reduced BTK levels and appeared to reduce the AAK1-annotated protein group (A0A096LP25), whereas NX-5948 reduced BTK alone (**A**, **B**). Western blotting confirmed AAK1 expression levels were not affected by NX-2127 (**Figure S2**). Neither PROTAC had a significant effect on the levels of TEC (**C**). The bar graph represents quantification of the western blots expressed as a percentage of the vehicle control (Mean ± SEM, n=3). NX-2127 degraded BTK by 94.3% while NX-5948 degraded BTK by 98.3% (**C**).

**Figure 3:**
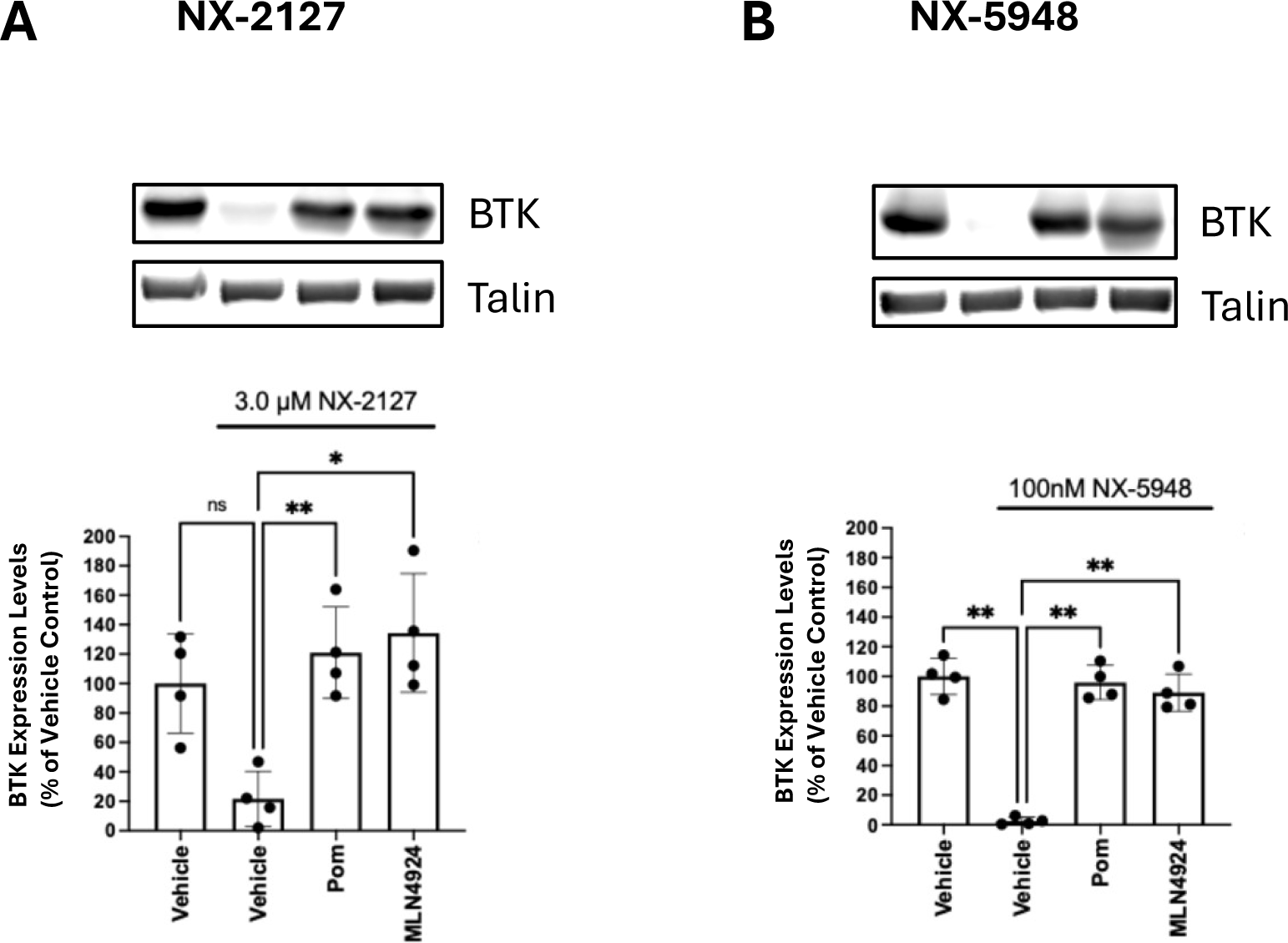
Effect of PROTAC Mechanistic Inhibitors. (A,B) Human platelets in PRP were preincubated with either vehicle (DMSO), 10 μM pomalidomide or 25 μM MLN4924 for one hour at 30°C. Following preincubation platelets were further treated with either 3 μM NX-2127 (**A**) or 100 nM NX-5948 (**B**) for 4 hours at 30°C. The graph represents quantification of the western blots expressed as a percentage of the vehicle control (Mean ± SEM, n=4). Representative blots are shown above the graph. Both PROTACs function as anticipated based on their mechanism of action. Pomalidomide, an IMiD-class CRBN binder, prevents BTK degradation by competing with both NX-2127 and NX-5948 for binding to the E3 ligase cereblon. MLN4924, a neddylation inhibitor of Cullin-RING ligases, blocks BTK degradation by inhibiting the transfer of ubiquitin, thereby preventing proteasomal targeting.

### BTK degradation abolishes GPVI- but not PAR-1-dependent Ca²⁺ signalling

As BTK operates downstream of ITAM-linked receptors such as GPVI, but not of G-protein-coupled receptors like PAR-1, we measured cytosolic Ca²⁺ mobilisation following BTK degradation. TRAP-6-induced PAR-1 signalling remained unchanged after treatment with either degrader (**Figure 4A**). In contrast, Ca²⁺ mobilisation in response to the GPVI agonist CRP was almost completely abolished across agonist concentrations up to 10 µg/ml, with only minimal residual flux at 20 µg/ml (Figure 4A). NX-5948 produced the most profound inhibition, achieving full suppression at doses that preserved overall platelet viability. These data support BTK as a non-redundant mediator of GPVI-mediated PLCγ₂ activation and Ca²⁺ release, but dispensable for PAR-1 signalling. The preservation of PAR-1 responses indicates that overall platelet viability and signalling competence are maintained.

**Figure 4:**
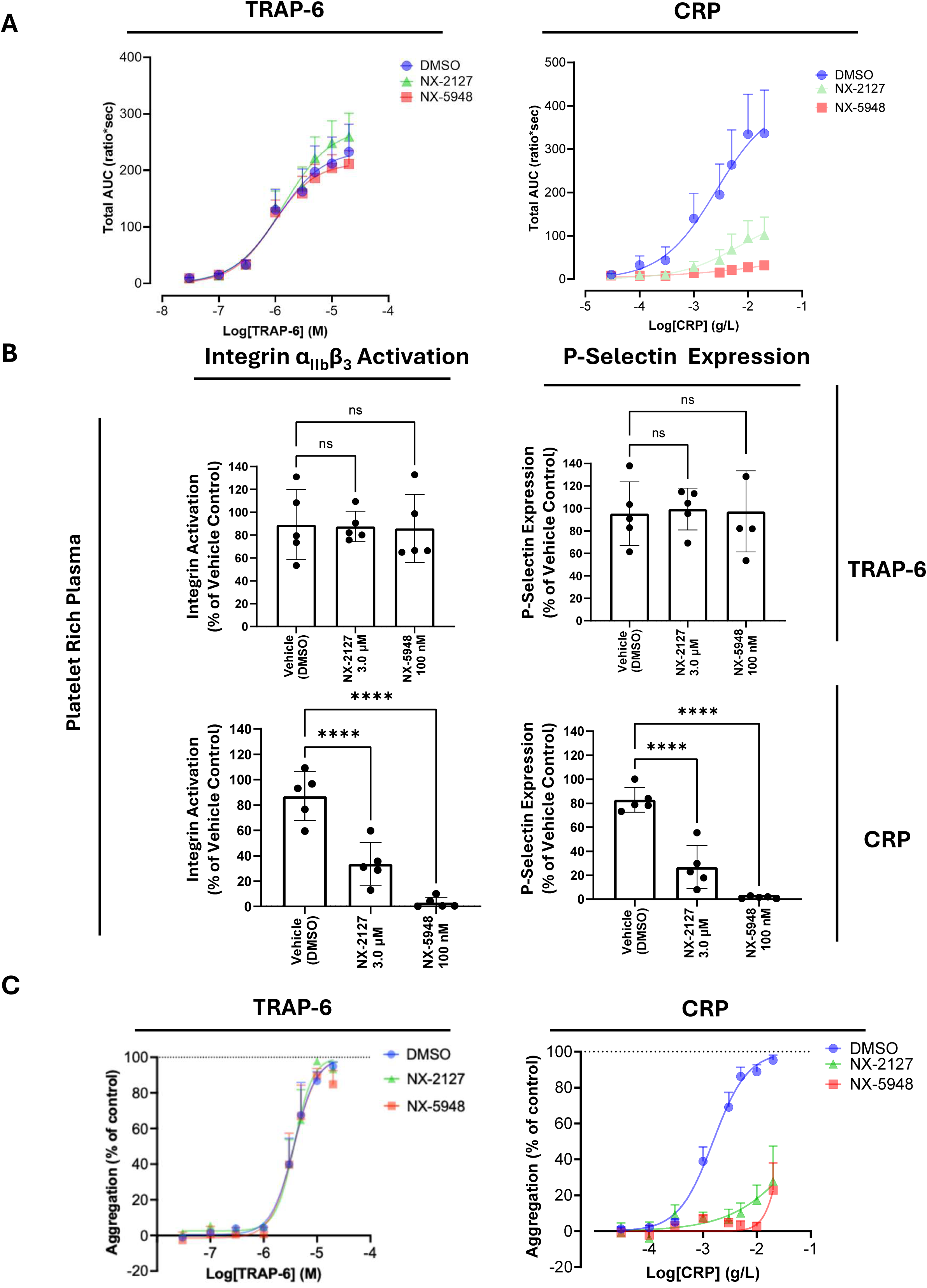
BTK Degradation impairs CRP-, but not TRAP-6-mediated platelet function. (A-C) Human platelets in PRP were treated with 3.0 μM NX-2127 and 100 nM NX-5948 for 4 hours at 30°C. Ca^2+^ mobilisation was assessed in Fura-2 loaded platelets in response to various concentrations of TRAP-6 and CRP (Data were expressed as the 340/380 nm fluorescence ratio) (**A**). Platelet activation was assessed by stimulation with either TRAP-6 or CRP then measuring integrin α_IIb_β_3_ activation and P-selectin expression by flow cytometry **(B)**. Data are expressed as a percentage of vehicle control (Mean ± SEM, n=5). There were no significant changes in platelet activation in NX-2127 or NX-5948 treated platelets following stimulation with TRAP-6. However, CRP-mediated platelet activation was significantly reduced by both NX-2127 and NX-5948 treatment, with NX-5948 having a stronger effect (**B**). Platelet aggregation (optical aggregometry) was unaffected in platelets stimulated by TRAP-6 following NX-2127 and NX-5948 treatment. However, BTK degradation by NX-2127 and NX-5948 severely impacted CRP-mediated platelet aggregation (**C**) Data are expressed as percentage of max response to platelet poor plasma (PPP)(Mean ± SEM, n=5).

### NX-5348 selectively suppresses GPVI-dependent activation and aggregation

Flow cytometric analysis revealed that BTK degradation translated directly into impaired functional activation. CRP-stimulated integrin α_IIb_β₃ activation and P-selectin expression were sharply reduced by both degraders, with NX-5948 producing near-complete inhibition (**Figure 4B**). In contrast, TRAP-6-induced responses were unaffected, confirming that PAR-1-mediated platelet function was intact. Aggregometry mirrored these results: CRP-induced aggregation was strongly suppressed, whereas thrombin and TRAP-6 responses were unchanged. These functional data show that NX-2127 and NX-5948 selectively impair GPVI-dependent, but not PAR-1-dependent, platelet activation.

### BTK loss arrests platelet spreading at the filopodia stage

As GPVI also contributes to platelet spreading on a collagen surface^26^, we assessed platelet adhesion and spreading under static conditions. Adhesion to collagen was preserved after NX-2127 or NX-5948 treatment, but there was a non-significant trend toward a reduction of the proportion of fully spread platelets, producing a morphology dominated by filopodia (**Figure 5A**). On fibrinogen, NX-5948 significantly reduced the lamellipodia-to-filopodia ratio (**Figure 5B**), consistent with impaired integrin α_IIb_β₃-dependent spreading. These findings show that BTK is not involved in initial adhesion of platelets but may promote progression to full spreading, including lamellipodia formation.

**Figure 5:**
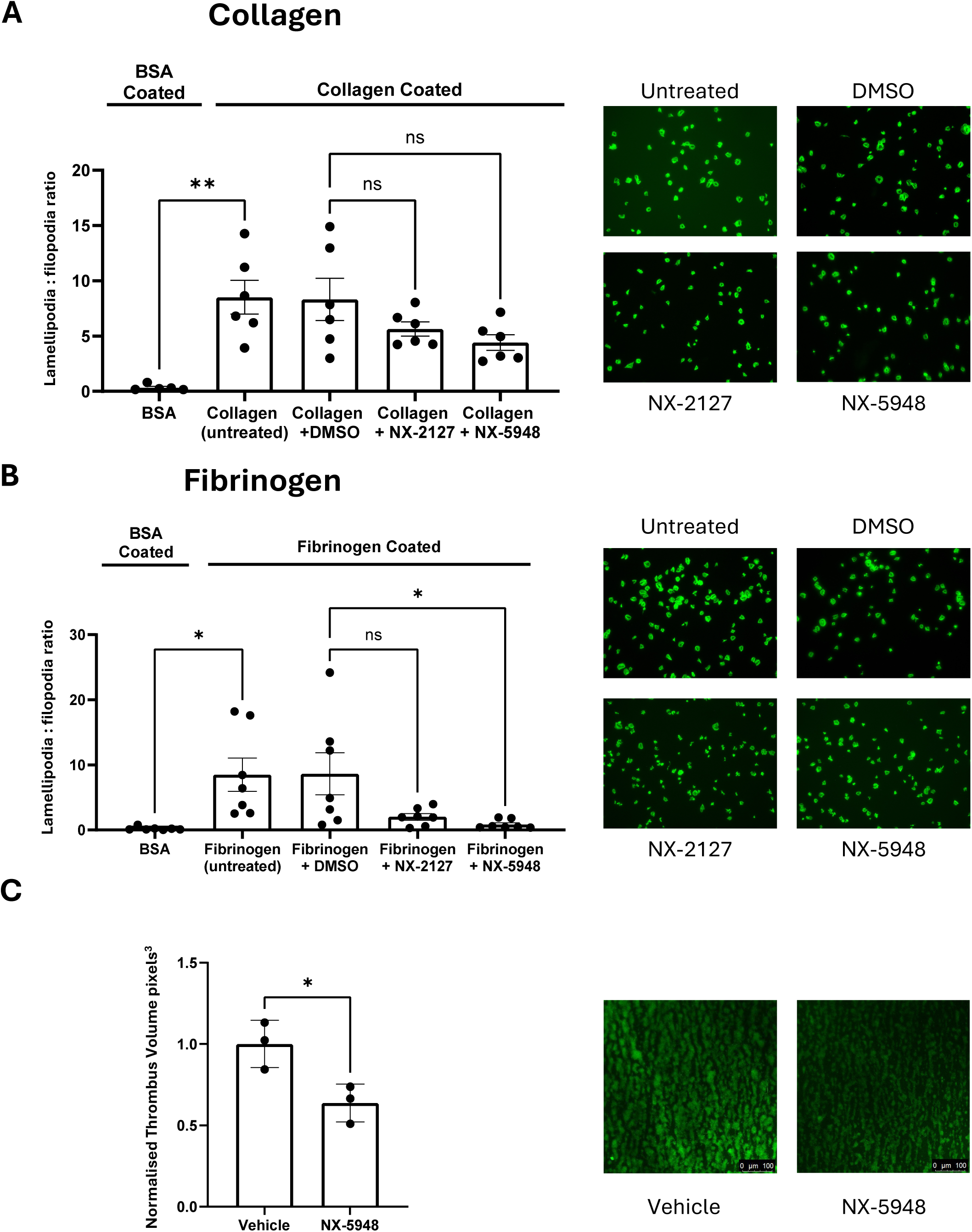
Effect of BTK Degradation on platelet spreading and thrombus formation. (A,B) Human platelets in PRP were treated with 3.0 μM NX-2127 and 100 nM NX-5948 for 4 hours at 30°C. Platelets were washed and added to surfaces coated with 10 µg/ml collagen **(A)** or 20 µg/ml fibrinogen **(B)** and allowed to adhere for 75 min. Platelets were stained with ActinGreen and visualised by confocal microscopy. Five images were taken per coverslip for each condition, and platelet spreading stage (round/filopodia/lamellipodia) was analysed using FIJI (ImageJ). The graphs show the lamellipodia/filopodia ratio for each condition. BSA represents spreading on BSA-coated coverslips (negative control), and “Collagen/Fibrinogen (untreated)” indicates platelets spread on extracellular matrix (ECM) without DMSO (positive control). (A: Mean ± SEM, n=6; B: Mean ± SEM, n=7) **(C)** Human platelets in PRP were treated with 100 nM NX-5948 for 4 hours at 30°C. Platelets were stained with DiOC6, recombined with the red blood cell layer and whole blood was passed over a 50 µg/ml collagen-coated surface under flow (3.07 ml/h (shear rate = 1000 s⁻¹) for 5 min). Representative images are shown on the right and thrombus volume on the left (Mean ± SEM, n=3).

### NX-5348 suppresses thrombus formation in vitro and in vivo while preserving haemostasis

Having established NX-5948 as a highly potent, fast, precise, and mechanism-validated BTK degrader in human platelets, we advanced NX-5948 to in vitro and in vivo thrombosis models. Under flow conditions on collagen, NX-5948 markedly reduced thrombus volume and stability (**Figure 5C**). In mice receiving a single 30 mg/kg oral dose, platelet Btk was reduced by 97% (**Figure 6A**). Ex vivo, GPVI-dependent integrin α_IIb_β₃ activation and P-selectin exposure were suppressed, whereas thrombin responses remained unaltered (**Figure 6B**).

**Figure 6:**
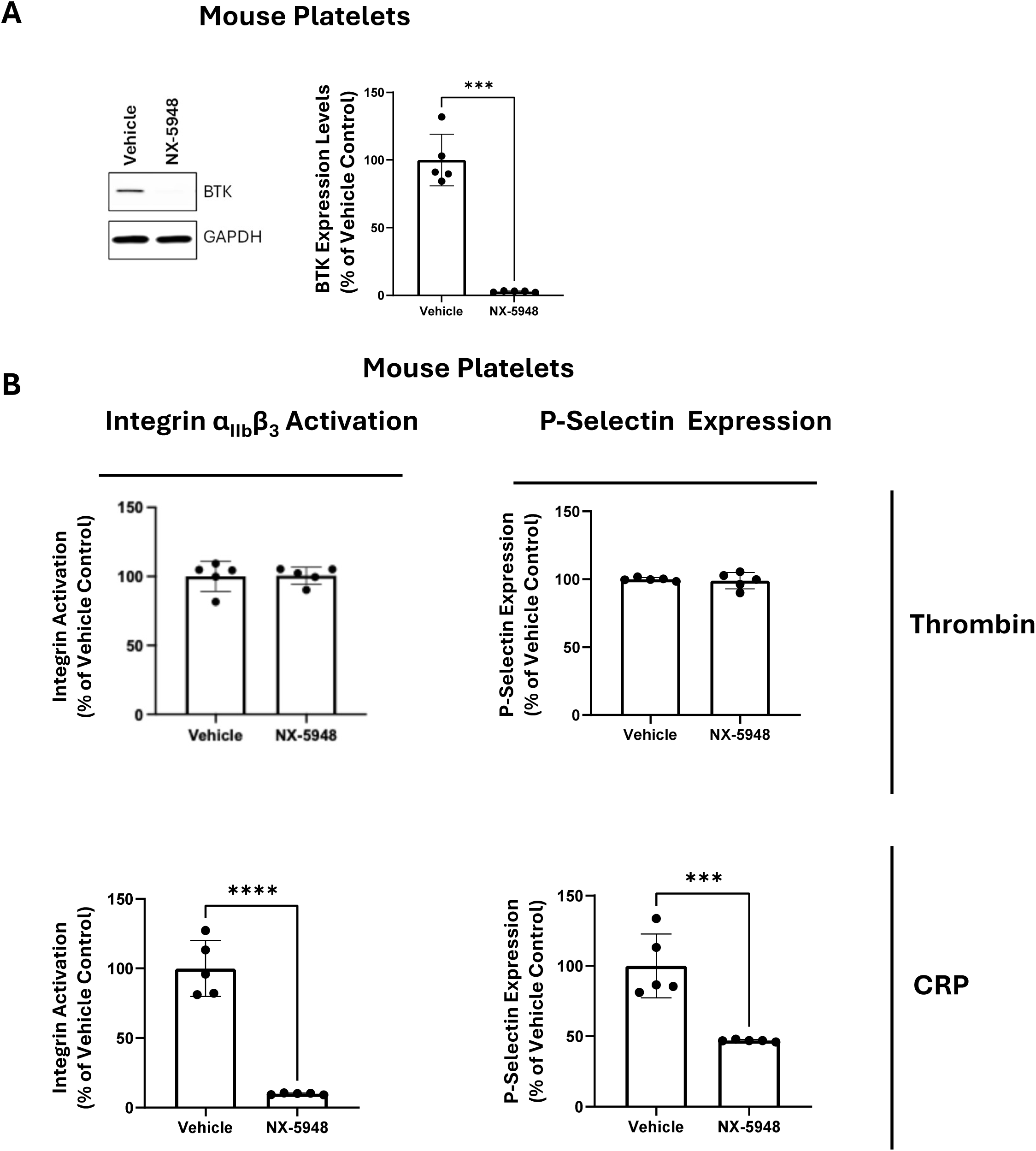
NX-5948 impairs *ex vivo* CRP-mediated platelet function in mice. (A,B) Mice were treated with 30 mg/kg of NX-5948 by oral gavage for 24 hours. Platelets were isolated and washed. Btk expression was measured by western blotting (**A**). The graph represents quantification of the western blots expressed as a percentage of the vehicle control (Mean ± SEM, n=5). Btk levels in mouse platelets were reduced by over 97% **(A)**. In platelets from mice treated with NX-5948, CRP-mediated platelet activation was significantly reduced when measured by both α_IIb_β_3_ integrin activation and P-selectin expression (CRP and thrombin used at 5 µg/ml and 1 U/ml respectively) (**B**). Thrombin mediated platelet activation was unaffected. Data are expressed as percentage of vehicle control (Mean ± SEM, n=5) **(B)**.

In vivo, NX-5948 treatment profoundly inhibited arterial thrombus formation following FeCl₃-induced carotid injury, reducing integrated fluorescence by > 70% relative to controls (**Figure 7A**). Importantly, tail-bleeding times were unaltered (**Figure 7B**), demonstrating that potent suppression of pathological thrombosis can be achieved without compromising haemostasis.

**Figure 7:**
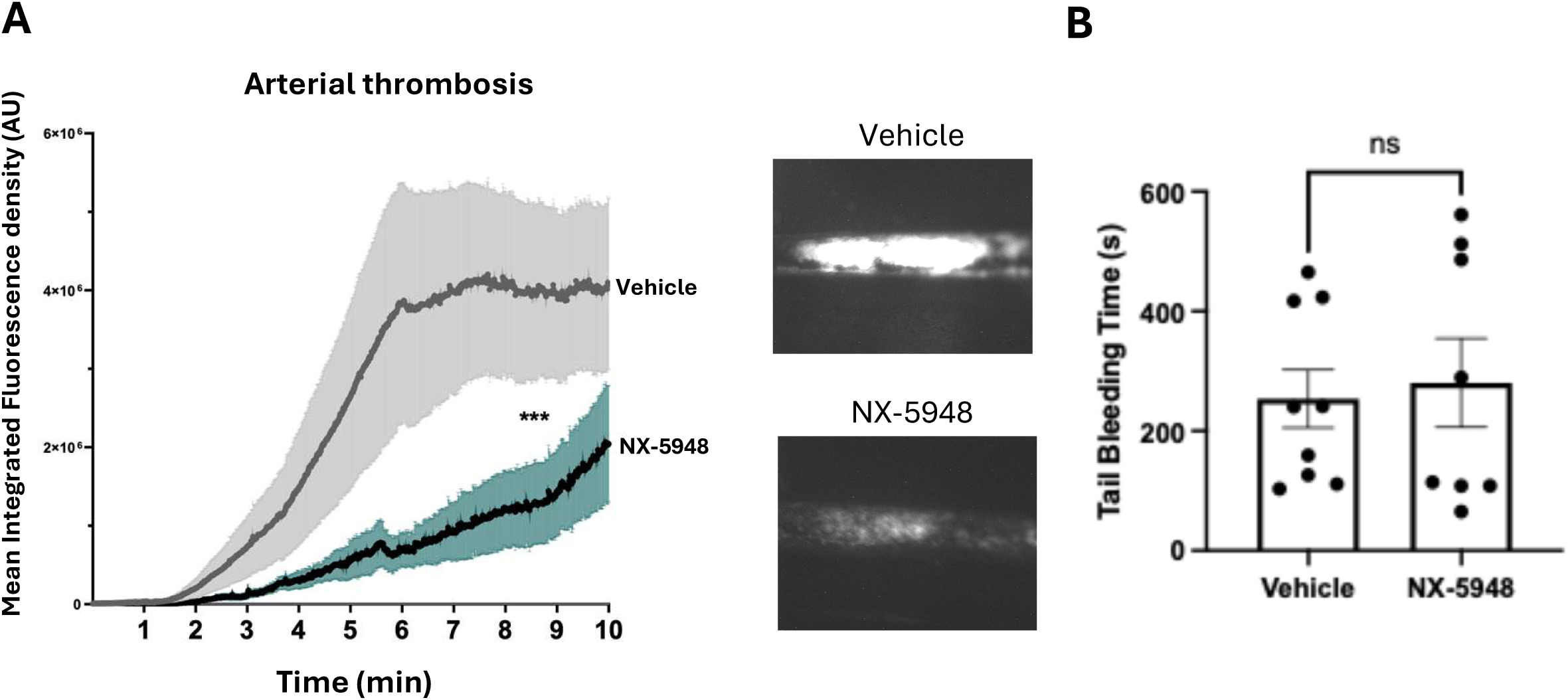
Impaired arterial thrombosis in NX-5948 treated mice. (A,B) Mice were treated for 24 hours with either vehicle (DMSO) or 30mg/kg NX-5948 by oral gavage. Mice were anaesthetised using ketamine. Ferric chloride was applied to the carotid artery of mice to inflict a vascular injury and induce thrombus formation (12% for 3 minutes) **(A)**. Thrombus formation was imaged by intravital microscopy in real time. The data are expressed as mean integrated fluorescence density of the thrombus (Mean ± SEM, n=6), indicative of thrombus size and density **(A)**. Tail bleeding was assessed following a tail incision. The data are expressed as bleeding time in seconds (Mean ± SEM, n=9). There is no significant difference between the bleeding time of vehicle and NX-5948 treated mice **(B)**.

Together, these results establish NX-5948 as a first-in-class, orally active BTK degrader that achieves rapid, complete, and selective BTK removal in platelets, silencing GPVI-dependent activation while preserving PAR-1-mediated function. This pharmacological separation between thrombosis and bleeding represents a major conceptual advance for antithrombotic therapy and validates targeted protein degradation as a viable strategy in anucleate cells.

## Discussion

BTK has emerged as a promising antithrombotic target, and here we compared the degraders NX-2127 and NX-5948 to define their effects on platelet BTK, function, and thrombosis. Both compounds efficiently degraded BTK in human and mouse platelets, but NX-5948 showed greater potency, faster kinetics, and deeper degradation, consistent with cooperative binding. Proteomics confirmed the high selectivity of both degraders. Functionally, both compounds suppressed CRP-induced integrin α_IIb_β₃ activation, P-selectin expression, and aggregation, but NX-5948 produced near-complete inhibition of GPVI signalling. TRAP-6-mediated PAR-1 signalling remained intact, confirming pathway specificity. In vivo, oral NX-5948 markedly reduced ferric-chloride-induced arterial thrombosis in mice without prolonging bleeding. Together, these data establish NX-5948 as a potent and selective BTK degrader that prevents thrombosis while preserving haemostasis, supporting its further development as a next-generation antithrombotic.

We previously demonstrated that BTK degraders such as DD-03-171 and DD-04-15 target platelet BTK but also reduce TEC by ∼75%^19^. In contrast, NX-2127 and NX-5948 showed minimal effects on TEC, providing more selective tools to dissect BTK function. Among these, NX-5948 was clearly superior, showing greater potency, more rapid kinetics and near complete BTK degradation. BTK, a non-receptor tyrosine kinase of the TEC family, mediates platelet activation downstream of GPVI, CLEC-2, and FcγRIIa through ITAM or hemITAM motifs. Following GPVI ligation, Lyn phosphorylates FcRγ, recruiting SYK and the LAT signalosome, where BTK becomes activated to phosphorylate PLCγ2 and drive Ca²⁺ mobilisation, key events in integrin α_IIb_β_3_ activation, secretion, and thrombus growth. Although TEC can compensate for BTK deficiency in mice^27^, our findings confirm BTK as the principal mediator of GPVI-dependent Ca²⁺ signalling, consistent with observations from XLA patients whose platelets fail to mobilise Ca²⁺ in response to CRP^15^.

Both degraders markedly inhibited CRP-mediated integrin α_IIb_β_3_ activation and aggregation, yet TRAP-6 responses were unaffected, reinforcing pathway specificity. Despite BTK’s key role in GPVI signalling, platelet adhesion to collagen under static or flow conditions remained intact, consistent with previous inhibitor studies^28,29^. While both compounds modestly reduced spreading on collagen, this did not reach significance. The absence of a clear adhesion defect may reflect residual GPVI-independent signalling or compensation by TEC, although the lack of effect of dual BTK/TEC inhibitors on adhesion and spreading would argue against this^28^. Other receptors such as integrin α_2_β_1_ and GPIb may also contribute under these conditions^27^. Integrin α_IIb_β_3_ -mediated outside-in signalling is also linked to PLCγ2 and calcium dynamics through BTK^27,30^. On fibrinogen, NX-5948 significantly reduced the filopodia-to-lamellipodia transition. Collectively, these data indicate that BTK can promote platelet spreading and cytoskeletal reorganisation but is not essential for adhesion, which may explain the preserved haemostatic function despite potent antithrombotic activity.

Given its superior pharmacological profile, NX-5948 was selected for in vivo evaluation. A single 30 mg/kg oral dose eliminated platelet BTK within 24 hours and significantly impaired GPVI-mediated activation and P-selectin expression ex vivo. Strikingly, NX-5948 markedly suppressed arterial thrombosis without prolonging bleeding, achieving a clear separation of antithrombotic efficacy from haemostatic safety.

BTK degradation may have additional therapeutic implications beyond thrombosis. Patients with B-cell malignancies, where NX-5948 is already in clinical development, exhibit elevated thrombotic risk, particularly for deep vein thrombosis (DVT). Experimental data identify platelet BTK as a key driver of DVT^12^, suggesting NX-5948 could confer dual benefit: antineoplastic and antithrombotic. A second potential indication is immune thrombocytopenic purpura (ITP), where BTK mediates both B-cell-driven autoantibody production and Fcγ-dependent platelet clearance. BTK inhibitors have shown efficacy in ITP^31–33^, and degraders such as NX-5948 could enhance this effect by simultaneously limiting antibody production, macrophage phagocytosis, and thrombotic risk, particularly in patients treated with thrombopoietin receptor agonists who experience paradoxical hypercoagulability^34^.

Because platelets cannot resynthesise proteins, BTK degradation will persist until new platelets are produced, suggesting an optimal dosing window that maintains antithrombotic efficacy while sparing other blood cells may exist. This pharmacological profile could benefit patients at high thrombotic risk who are intolerant of dual antiplatelet therapy (DAPT) due to bleeding. By reducing thrombus formation without impairing haemostasis, NX-5948 addresses a central limitation of current antiplatelet strategies^35^.

In summary, NX-5948 is a potent and selective BTK degrader that suppresses GPVI-but not PAR-1-mediated platelet activation, effectively limiting thrombosis without prolonging bleeding. Its mechanism of action, safety profile, and oral bioavailability highlight its potential as a next-generation antithrombotic that dissociates efficacy from bleeding risk and may offer additional clinical benefit in thrombo-inflammatory and autoimmune disease. By separating thrombosis from haemostasis via degradation, rather than inhibition, of BTK, NX-5948 defines a mechanistically precise antiplatelet strategy. These data validate targeted protein degradation in platelets and motivate formal dose-exposure-response and durability studies toward first-in-class antithrombotic development.

## Acknowledgements

This study was supported by the BBSRC; BB/X017176/1 (CJM, JT), BHF; FS/4yPhD/F/21/34162 (CW), SP/F/21/150023 (YL) and PG/21/10760 (JV), NC3R (JST), Bristol Alumni and Tocris Biotechne (HW).

## Authorship Contributions

Conceptualisation, CJM, JST and IH; Data Curation, CJM; Formal analysis, CJM, JST, CW, YL, JV, HW, JT; Funding acquisition, IH, AWP; Investigation, CJM, JST, CW, YL, JV, HW; Supervision, IH; Writing-original draft preparation, CJM; Writing-review and editing, IH.

## Authorship and conflict-of-interest statements

All authors approved the final version of the manuscript. The authors declare that they have no competing financial interests or conflicts of interest related to the work described in this article.

---

**Figure S1:** Concentration-dependent reduction of BTK expression by NX-5948 in whole blood. Concentration-degradation curve of NX-5948 in whole human blood. Whole blood was treated with the indicated concentrations of NX-5948 for 4 hours at 30°C. BTK levels were quantified by densitometry of western blot and normalised to vehicle (DMSO) treated control. The curves were generated from quantification of western blots (Mean ± SEM, n=3 with representative blots shown above the respective curve. The degradation curve for NX-5948 in whole blood was similar for those in Figure 1D, suggesting limited sequestration by other blood cells.

**Figure S2:** Concentration-dependent reduction of AAK1 expression by NX-5948 in PRP. Human platelets in PRP were treated with 3.0 μM NX-2127 for 4 hours at 30°C. Platelets were washed and BTK and AAK1 levels were quantified by densitometry of western blot and normalised to vehicle (DMSO) treated control (Mean ± SEM, n=3).

