## Supplementary figures and images for "Selective degradation of platelet BTK by PROTAC NX-5948 provides antithrombotic benefits without affecting haemostasis"

### Supplemental Figures S1 and S2

#### Figure S1

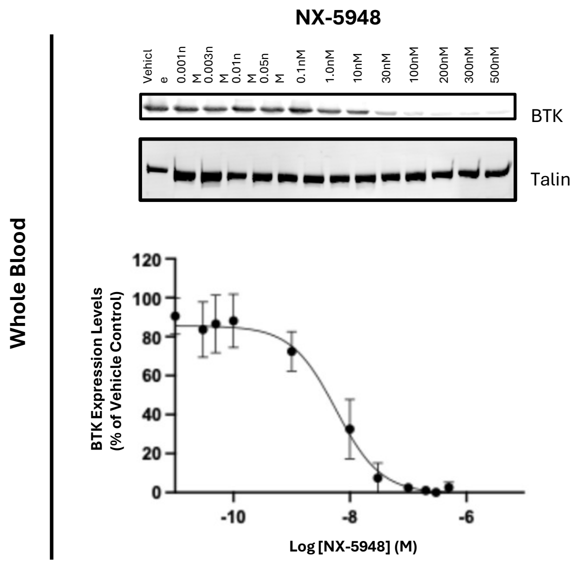


#### Figure S2

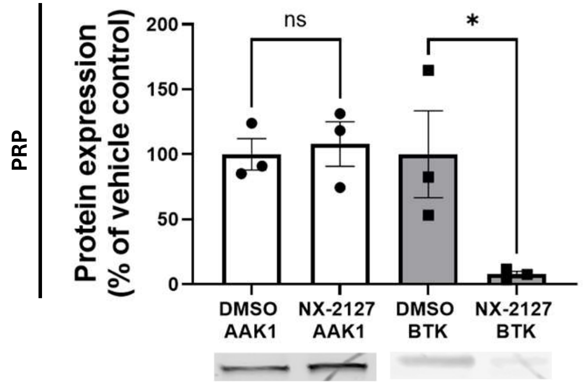
